# Transfer Learning for Inference of Metastatic Origin from Whole Slide Histology

**DOI:** 10.1101/2021.04.21.440864

**Authors:** Geoffrey F. Schau, Hassan Ghani, Erik A. Burlingame, Guillaume Thibault, Joe W. Gray, Christopher Corless, Young Hwan Chang

## Abstract

Accurate diagnosis of metastatic cancer is essential for prescribing optimal control strategies to halt further spread of metastasizing disease. While pathological inspection aided by immunohistochemistry staining provides a valuable gold standard for clinical diagnostics, deep learning methods have emerged as powerful tools for identifying clinically relevant features of whole slide histology relevant to a tumor’s metastatic origin. Although deep learning models require significant training data to learn effectively, transfer learning paradigms provide mechanisms to circumvent limited training data by first training a model on related data prior to fine-tuning on smaller data sets of interest. In this work we propose a transfer learning approach that trains a convolutional neural network to infer the metastatic origin of tumor tissue from whole slide images of hematoxylin and eosin (H&E) stained tissue sections and illustrate the advantages of pre-training network on whole slide images of primary tumor morphology. We further characterize statistical dissimilarity between primary and metastatic tumors of various indications on patch-level images to highlight limitations of our indication-specific transfer learning approach. Using a primary-to-metastatic transfer learning approach, we achieved mean class-specific areas under receiver operator characteristics curve (AUROC) of 0.779, which outperformed comparable models trained on only images of primary tumor (mean AUROC of 0.691) or trained on only images of metastatic tumor (mean AUROC of 0.675), supporting the use of large scale primary tumor imaging data in developing computer vision models to characterize metastatic origin of tumor lesions.

## 1 Introduction

Cancers that spread to distal regions of the body are referred to as metastatic cancers, and often present a significant adverse clinical milestone of cancer evolution that accounts for a majority of deaths associated with solid tumors[1]. By invading into nearby tissue, or navigating transportation systems such as lymph or blood circulation, metastatic cancers cells migrate to new locations throughout the body where they can establish new residence and continue to proliferate. Common sites for cancer metastasis are the lung, bone, brain, and liver, each of which presents distinct microenvironmental conditions and factors that may affect the cancer cell’s capacity to divide and spread further[2]. Because cancer cells that have metastasized retain capacity for dislocation and traversal, and because many different cancer types can metastasize to the same site, accurate inference and diagnosis of metastatic cancer is essential for devising treatment strategies to control and halt further metastatic processes.

Generally, cancers that metastasize from one site, such as the colon, into a new site, such as the liver, retain similar morphological and structural features. Seen under a microscope, a colon cancer that has metastasized to the liver will generally appear more similar to a primary colon cancer than to a liver cancer that has yet to metastasize. Clinically, pathologists rely on visual inspection of H&E stained section of metastatic tumor tissue to infer the cancer’s origin. However, in challenging cases where a metastatic diagnosis is not readily obvious, confirmatory assays such as immunohistochemistry (IHC) staining can provide definitive diagnosis with which to guide treatment.

We previously demonstrated a learning system trained to classify whole slide images of cancers that had metastasized to the liver according to the tumor’s tissue of origin[3]. That work was limited to three of the most commonly recurring classes of metastatic cancer due in part to limited data availability of other classes of metastatic cancers. This work seeks to leverage morphological and spatial properties of primary tumor tissue to enhance the performance of a classification model trained to infer the origin of secondary metastatic cancer based on histopathological presentation in digital whole slide images. Similarities between primary and metastatic tumors have shown striking similarity in gene expression[4], growth characteristics[5], and chromosomal rearrangement[6] and more recent work has leveraged passenger mutations to accurately classify primary and metastatic cancer based on genomic signatures [7]. Motivated by primary-metastatic structural similarity evident in whole slide histology, this work seeks to evaluate whether a computer vision system trained to classify primary cancers retains predictive power when inferring the origin of metastatic cancer based on similarities in tissue morphology.

Training a model in one setting and transferring it into a different setting is an example of a transfer learning paradigm. These approaches have demonstrated robust capacity for boosting model performance[8], and have found wide use in the field of deep learning for computer vision applications[9, 10]. This work extends previous analyses by evaluating the degree to which a computer vision model generalizes to unseen samples of whole slide metastatic cancer by training on only primary cancers, only metastatic cancers, and by first training on primary and transferring the learning model to retrain on metastatic samples. Further, this work evaluates the divergence between primary and metastatic cancers of different types within learned unsupervised morphological feature space, and draws connections between the degree to which primary and metastatic cancers are dissimilar and how well models generalize to correctly predicting metastatic origin of different cancer types.

## 2 Materials and Methods

An overview of the computational pipeline developed in this work is shown in Figure 1. Our approach separates the study objectives into two components. The first portion of the study pre-trains a neural network classifier to predict tumor type from whole slide images of primary cancers, while the second portion transfers the learned classification model from primary cancers into a data set composed of metastatic tumors to infer the samples’ metastatic origin. To evaluate the efficacy of a primary-to-metastatic learning paradigm, we compare metastatic classification performance from three models, a first model trained only on images of primary tumor, a second model trained only on images of metastatic tumor, and a third model first trained on primary cancers that is then transferred to and fine-tuned within the metastatic cancer setting.

**Figure 1:**
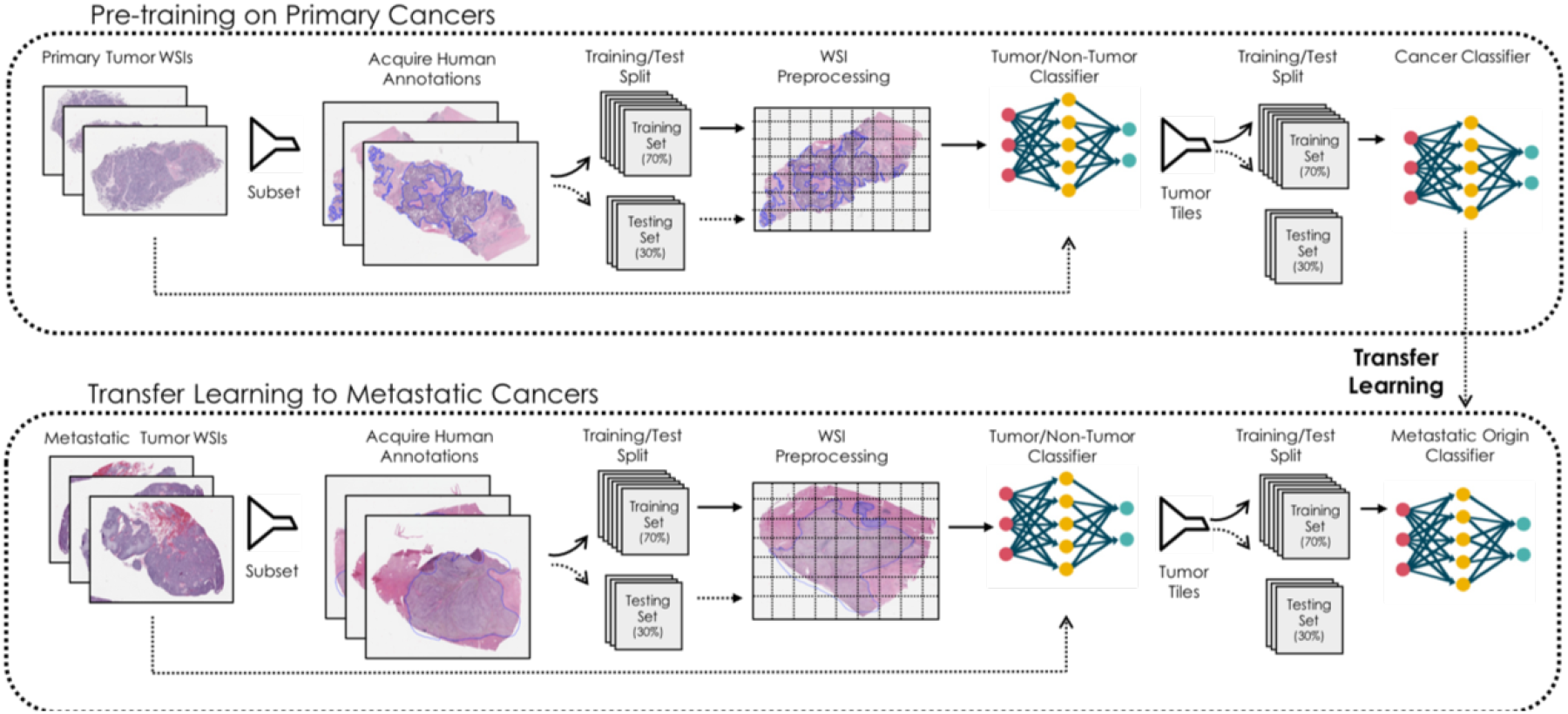
Transfer learning concept designed to leverage morphological and spatial features of primary cancer to infer the metastatic origin of secondary cancers. Whole slide images are tiled into non-overlapping square patches of 128 pixels at 20x magnification. In the top diagram, a subset of whole slide images are annotated by an expert pathologist to identify tumor tissue in the slide. These annotations are then employed to train a first-stage classification model that identifies tiles containing tumor tissue. The filter model is applied to the remaining data to identify tumor tissue within each of the whole slide images. Tiles classified as tumor tissue are then passed through a second model trained to correctly classify primary tumor type based on clinical annotation of the whole slide. In the bottom diagram, a similar first-stage classification model is trained to filter normal tissue surrounding the metastatic cancer prior to a second-stage model trained to correctly classify the metastatic origin of the tumor tissue. We evaluate the efficacy of transfer learning by fine-tuning a learning model trained on primary tumors with images of metastatic cancers.

Because whole slide images are large and heterogeneous, they often contain both tumor and non-tumor tissue. This mixture of tissue types has been shown to confound the degree to which classification models are generalizable, and so both components of this study employ a model to first identify tumor tissue from non-tumor tissue within the whole slide image trained on manual annotation of whole slide images by a board-certified pathologist. Annotated whole slide images are divided into training and test sets from which a ResNet50[11] deep learning model was trained to generate binary classifications of tumor and non-tumor tissue at patch-level resolution. These preliminary filter models are deployed onto their respective whole data sets to filter out normal stroma tissue from the tiled data sets such that the resultant data sets are composed of tumor tissue. Secondary models are then trained to correctly classify the tumor tiles according to their tissue of origin as informed by the clinical diagnostic record. Generally, metastatic cases are differently prevalent in clinical records, resulting in imbalanced representations of metastatic classes. In this work, samples were collected from fourteen common tumor types from both primary and metastatic cancers that metastasize to the liver, as shown in Figure 2A.

**Figure 2:**
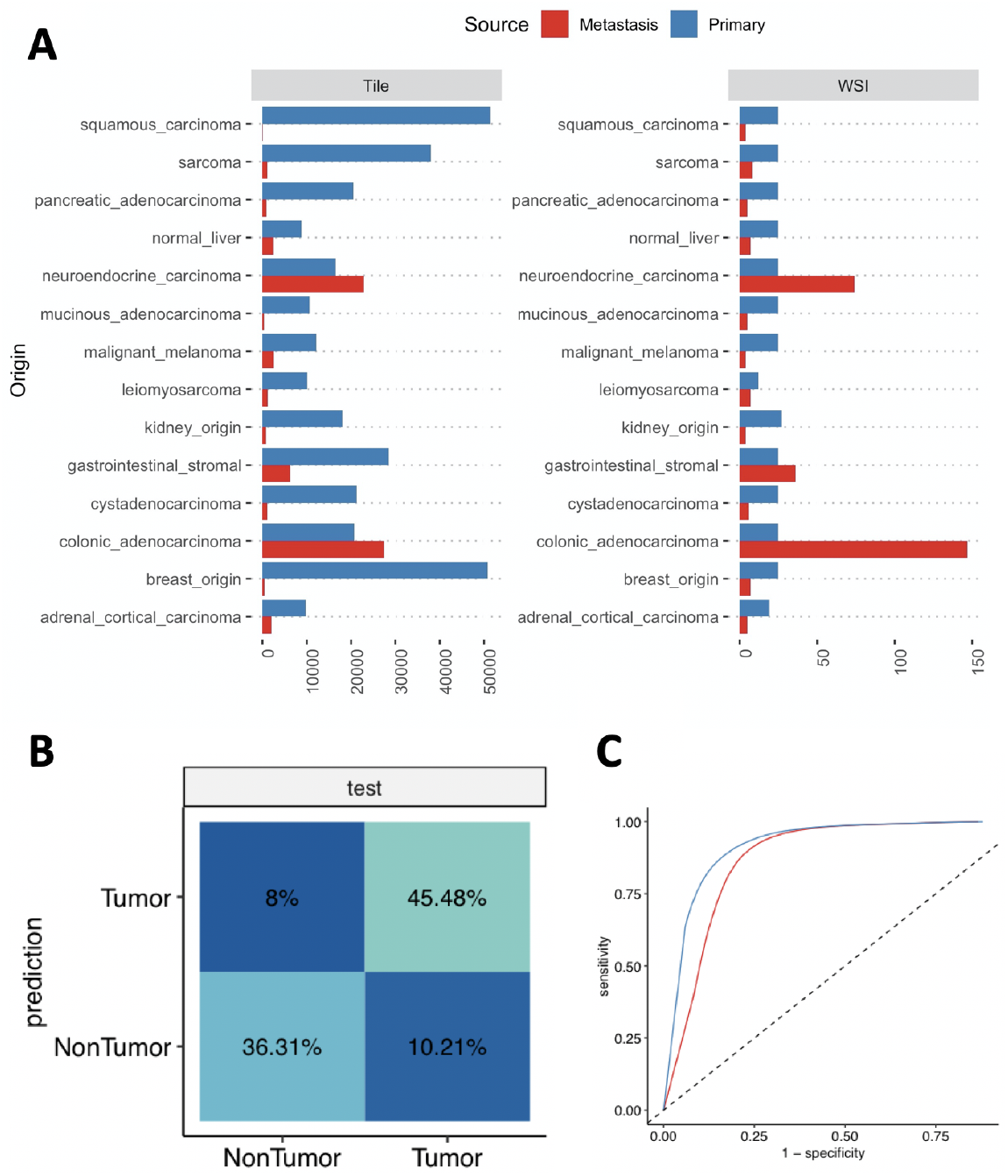
(A) Data used in this study for both primary and metastatic examples. Due to limitations in data availability for rare metastatic cancers, data is imbalanced in both whole slide and tile bases. To evaluate the efficacy of a primary-to-metastatic transfer learning paradigm, we explicitly balance a dataset of primary tumors to contain the same number of whole slide images. Variation in tile counts is due to natural variation in histological specimen size. (B) First-stage tumor filtering classification model performance on held-out testing data (red line, AUROC of 0.88) is similar to performance on seen training data (blue line, AUROC 0.92). (C) Confusion matrix of tumor filter model trained on pathological annotation of tumor regions within whole slide images.

### 2.1 Data Acquisition and Pre-processing

This work employs a dataset composed of 324 whole slide images of metastatic cancers and 344 whole slide images of primary cancer collected from the Knight BioLibrary and Knight Diagnostic Laboratories at Oregon Health & Science University (OHSU). Each whole slide image is divided into non-overlapping tiles 128 × 128 × 3 pixels wide that cover 100 *μ*m square. Tiles containing predominantly white background light were filtered out and the remaining tiles were color-normalized to a reference staining density[12]. Annotation tables associate each whole slide image with its tissue of origin informed by the clinical diagnostic record which are employed as the target variables for the classification model. In this work, while predictions are made on per-tile basis by the learning model, whole slide predictions are made as the mean prediction of each slide’s constituent tiles. The inherent class imbalance in the metastatic whole slide training set presented a limiting factor for previous work [3], which was limited to three principal classes of metastatic origin. This challenge is intended to be overcome in part by a transfer learning paradigm by which a larger related data set may be utilized to overcome class imbalance limitations for rare cases in metastatic cancers.

Annotations of tumor regions are used to train a tumor identification model and were drawn by an expert pathologist using the QuPath[13] software tool for 78 whole slide images spanning all fourteen distinct primary tissue types. Annotations were computationally extracted and employed to label each tile sampled from each of the 78 whole slide images as either belonging to a region annotated as tumor or not, in which case the tile is labeled non-tumor. The tumor-identification model is similar to previously-described work [3] in which a convolutional neural network was trained to identify tumor regions based on pathologist annotation.

### 2.2 Learning System Architecture

This study seeks to evaluate whether transfer learning from primary cancer cases into the metastatic cancer classification setting confers a computational advantage compared to primary- or metastatic-specific training. Here we utilize the ResNet50 learning architecture for both tumor region identification and tumor type classification. ResNet50 is a widely-used convolutional neural network based learning architecture that has been broadly applied to challenges in digital pathology[14, 15, 16]. We adopted the ResNet50 model and modified its output layer to contain fourteen nodes, each corresponding to one of the fourteen sites of tissue origin, and a softmax activation function such that the vector of output values sums to one, thereby enabling a probabilistic interpretation of the model’s output. We trained a single ResNet50 model for 10 epochs with a learning rate of 0.001 and batch size of 32 on the training set of whole slide image tiles in parallel on four Nvidia V100 GPUs for a total of 80000 weight update steps. Data loaders were designed to generate batches of data with the same data transforms that include flipping, rotating, and color scaling by saturation, brightness, and hue with class-balancing. In all cases, the Adam optimizer [17] was employed with a learning rate scheduler designed to decimate the learning rate at the end of each epoch. Data loaders were specifically designed to balance class representation to maximize class diversity with each batch of training data. To evaluate the transfer learning approach, a third model is trained for 5 epochs on primary cancer and fine-tuned for 5 epochs on images of metastatic cancer.

## 3 Results

This first-stage model designed to identify tumor tissue within whole slide histology is evaluated on a held-out test set and achieves an area under the receiver operator characteristics curve (AUROC) of 0.88, which is similar to the AUROC of 0.92 achieved on training data, shown in Figure 2B and 2C. Using this model as a first-stage tumor filter, the remaining tiles are passed through for classification according to the three approaches described above. The receiver operator characteristic curves for each of the three modeling approaches for correctly classifying metastatic origin of whole slide histologies are shown in Figure 3. In general, we observe a consistent improvement in classification when employing the transfer learning approach, shown in green, particularly in cases of cystadenocarcinomas, sarcomas, and colonic adenocarcinomas. However, we also observe a few instances where transfer learning appears to confer a negative effect on model performance, such as in the case of adrenal cortical carcinomas, gastrointestinal stromal tumors, and tumors of kidney origin. Underlying explanations for drop in performance using transfer learning for these cases are not fully understood, though performance differentials may be at least partially explained by indication-specific morphological heterogeneity and limited available training data to adequately capture all degrees of spatial variation. Nevertheless, we report a mean AUROC for the primary-to-metastatic transfer learning model of 0.779, which outperforms a similar model trained only on primary tumors (mean AUROC of 0.691) and a similar model trained only on metastatic tumors (mean AUROC of 0.675).

**Figure 3:**
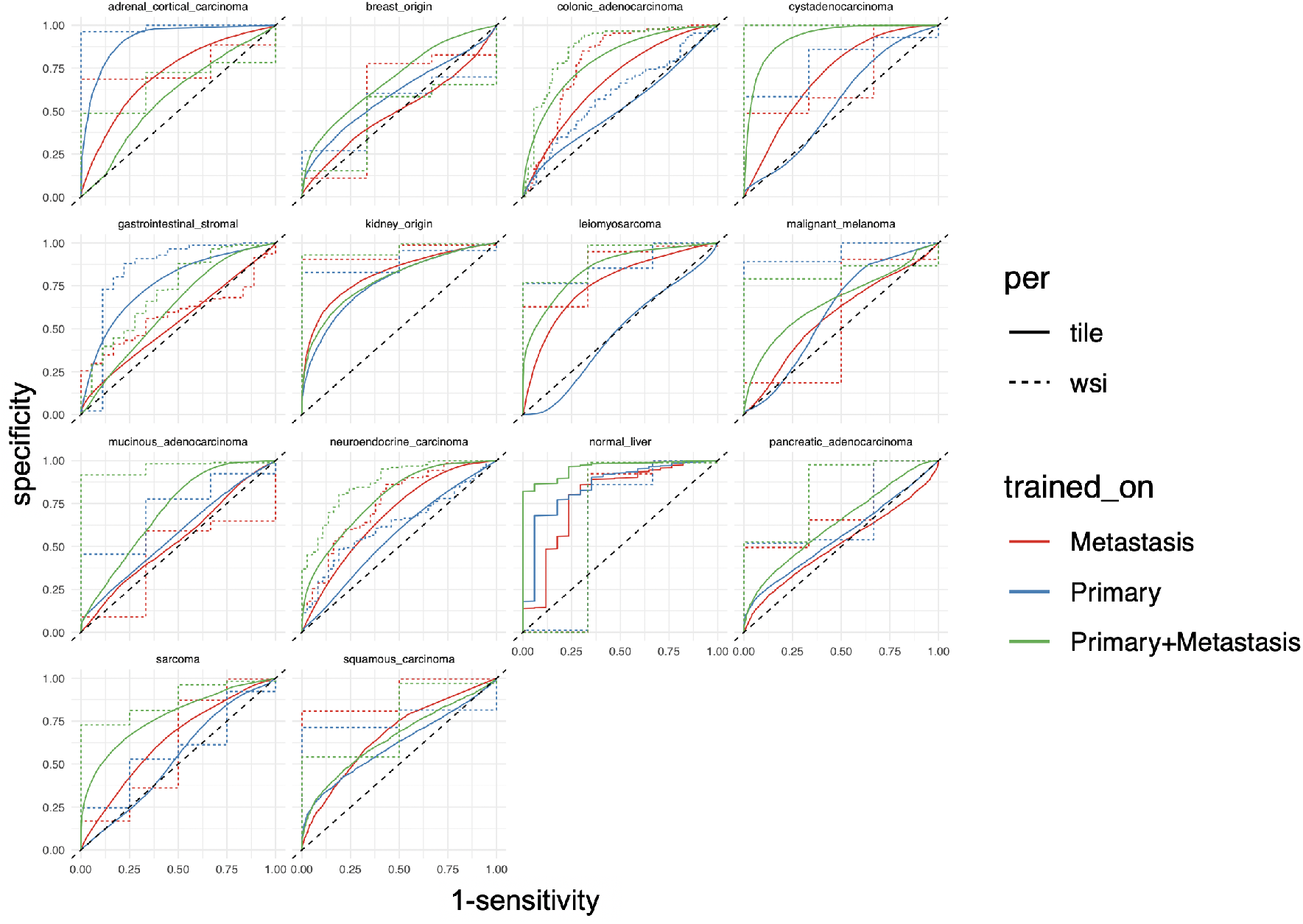
Receiver Operator Characteristics (ROC) curves shown for each of the fourteen class predictions by each of the three training strategies described above: exclusively trained on primary tumors, exclusively trained on metastatic tumors, and the proposed primary-to-metastasis transfer learning approach. In general, transfer learning confers a clear benefit to the model’s class-specific performance. However, adrenal cortical carcinomas and gastrointestinal stromal tumors appear to be exceptions in which transfer learning may adversely affect model performance. In each case we stratify the classification by tile (solid line) and by whole slide image (dashed line)

### 3.1 Spatial and Differential Predictions

The ROCs shown in Figure 3 evaluate classification performance for each class with respect to each of the remaining classes, providing a metric for how often the learned model’s maximum likelihood prediction correctly aligns with the true metastatic origin associated with each sample. Here we evaluate both spatially-refined predictions as well as the differential diagnoses generated by the best-performing model. Figure 4A illustrates how per-tile predictions are distributed within an example whole slide image. Class-confusion matrices are shown in Figure 4B which illustrate the degree to which certain cases are mistaken for others. In this example case shown in Figure 4A, the clinical diagnosis is a squamous carcinoma tumor, yet the model incorrectly predicted the case to be a gastrointestinal stromal tumor with 35% confidence. However, squamous carcinoma was the second most likely class with a model confidence of 18.3%. This perspective reflects that of a clinician’s differential diagnosis which may consider a set of underlying conditions responsible for manifestation of disease. We interpret the model’s differential diagnoses as the most likely predictions apart from the most likely prediction, analogous to how a practicing pathologist may suggest several potential indications which require confirmation through secondary assay such as an immunohistochemistry stain. Figure 4C illustrates this differential diagnosis for the trained model by plotting the rank of the true label on the x-axis. In the example shown in Figure 4A, the rank of the true label would be two, since the correct label was the model’s second most-likely prediction and the model’s confidence in that prediction on the y-axis. In this analysis we observe a steep roll-off as we move to the right, suggesting that although the model’s most likely prediction is correct 55% of the time, the probability that the correct guess is within the top six predictions from fourteen tumor types is greater than 90%.

**Figure 4:**
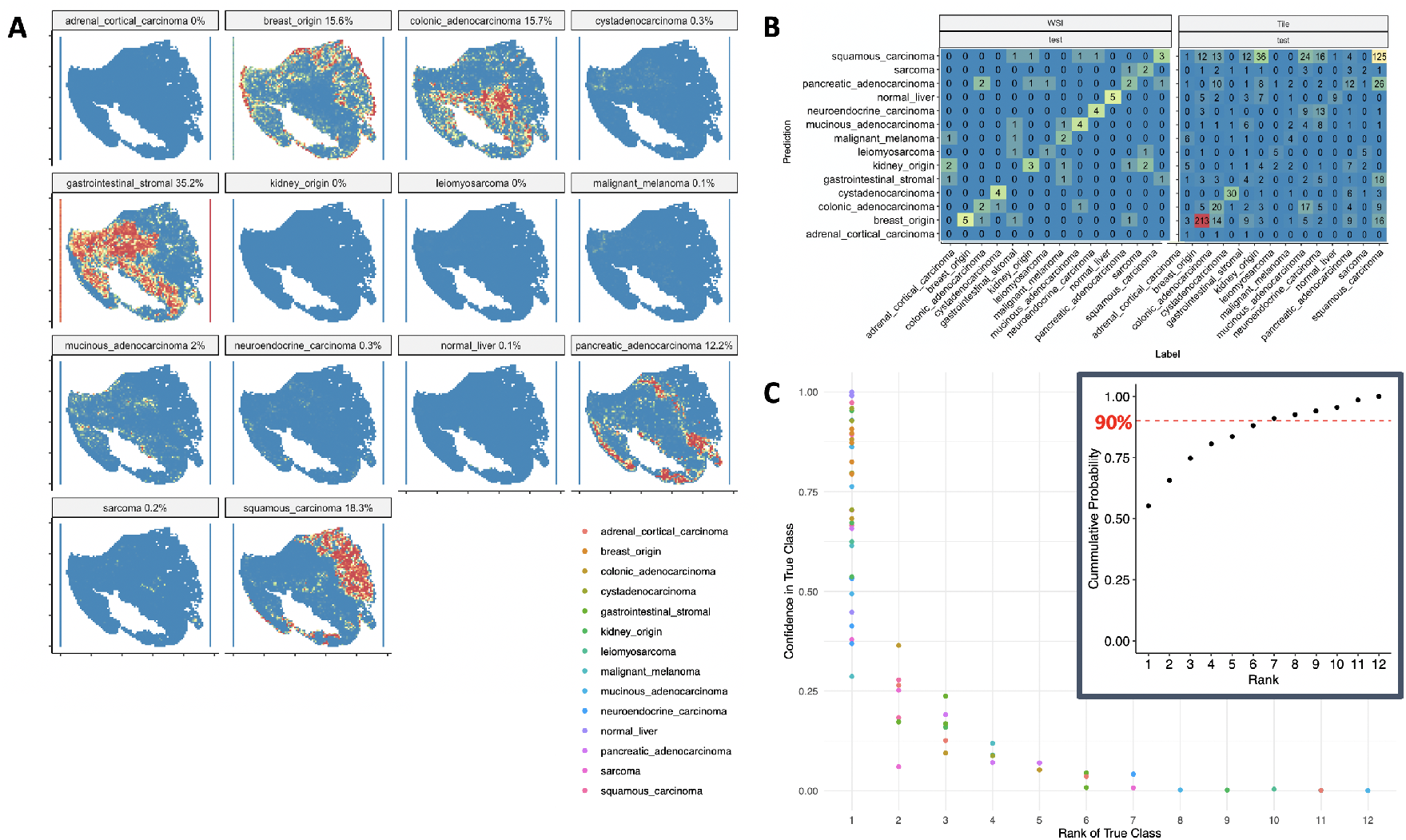
(A) Example heatmap showing spatially-resolved class-specific predictions made on each tile. This particular case shows a squamous carcinoma tumor, yet was incorrectly predicted to be a gastrointestinal stromal tumor by our model with 35% confidence. However, squamous carcinoma was the second most likely class with model confidence of 18.3%. (B) Confusion matrix from the held-out test set illustrating improved performance on whole slide images with respect to per-tile predictions. (C) Rank-confidence plot illustrating the degree to which the model confuses the correct prediction. A steep roll-off is reflected in the inset figure which measures the cumulative distribution of the rank-confidence values and suggests that while the model might not correctly predict the true class as its first choice, the correct choice is likely to be within the top six predictions with 90% likelihood.

### 3.2 Primary-Metastatic Feature Divergence

We next test the hypothesis that structural or morphological differences between primary tumors and their metastatic counterparts may affect the efficacy of our transfer learning approach. In this analysis, we randomly sub-sampled 5000 tiles from each of the 14 classes across each of the whole slide images in the study. As a sanity check, we include all tissue samples from normal liver from both primary and metastatic cancers, but otherwise filter out normal tissue from other origins using the classification models described above. With these tiles we train a 64-feature variational autoencoder (VAE) [18] to learn a latent representation of each of the histopathological tiles included in the entire data set used in this study. Latent representations are embedded into two dimensions using the t-stochastic neighbor embedding algorithm [19] and shown in Figure 5A. The 2D coordinate tSNE plane is faceted according to the densities of tiles drawn from each of the fourteen tumor types being classified and colored according to whether the tiles are drawn from images of either primary or metastatic tumor tissue. Intuitively, two distributions that are identical, meaning they share the same morphological features and image content, should perfectly overlap with each other, such as the instances of normal liver, which are expected to be highly concordant between primary and metastatic cancers. In principle, if tiles of a given cancer type look identical in both primary and metastatic images, then their distributions of tiles should perfectly overlap in this feature representation. Other examples show similar but not complete overlap, particularly in the cases of colonic adenocarcinomas and pancreatic adenocarcinomas. However, other samples appear to be more widely distributed and non-overlapping, such as neuroendocrine carcinomas and squamous carcinomas. Some cases, such as leiomyosarcomas, appear strongly bimodal, suggesting that some tiles share similar features while others do not.

**Figure 5:**
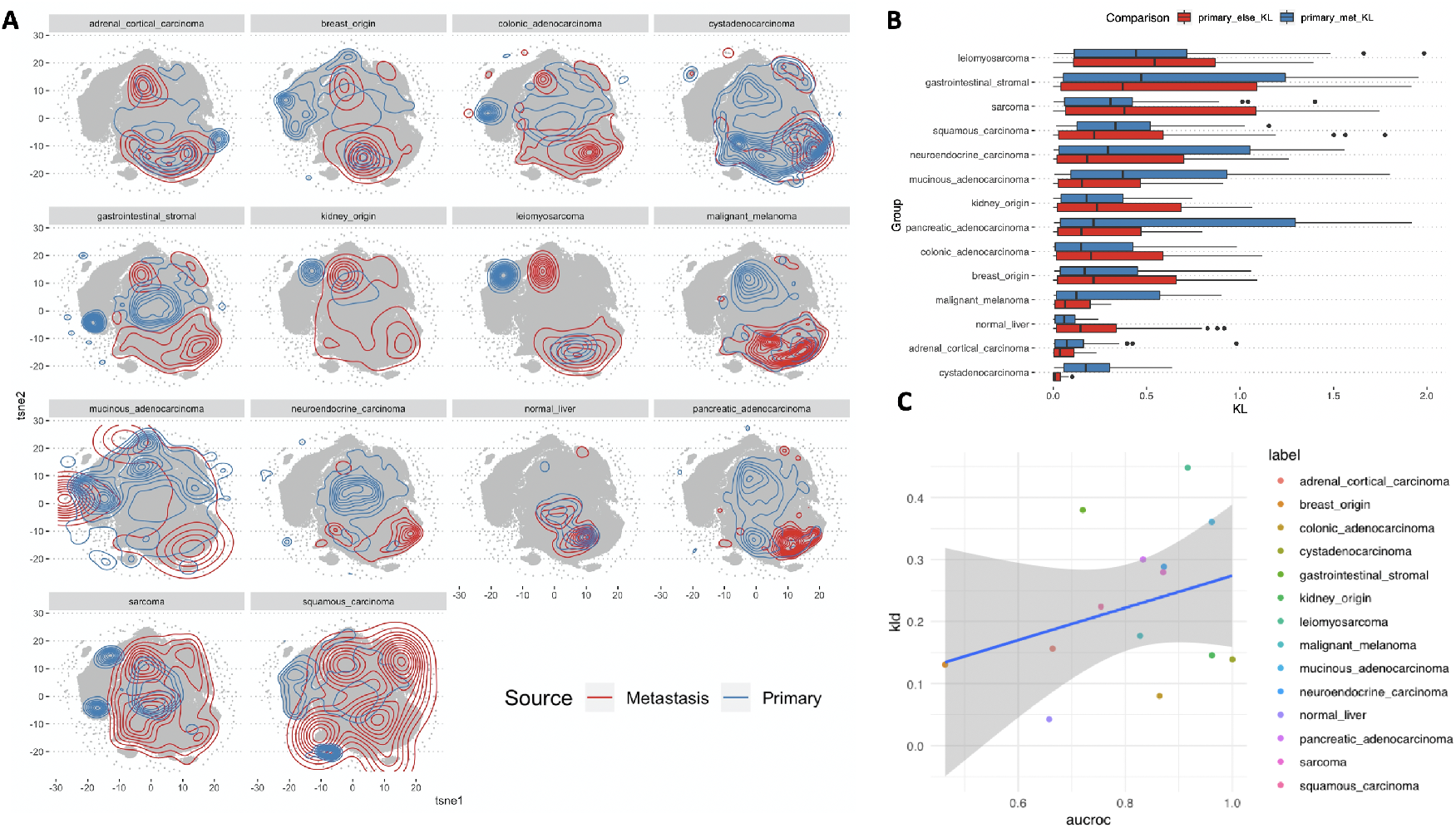
(A) A tSNE projection of image patches projected into two dimensions based on a learned representation of image content. Densities of tiles are shown for both primary and metastatic cases for each of the fourteen tumor types. Intuitively, if two distributions were perfectly overlapping, then their image content and tissue morphology distributions would be identical from images of primary and images of metastatic cancers. However, distributions that differ suggest shifts in morphology or image content following metastasis. (B) Quantitative estimation of distributional separation is computing using the Kullback-Leibler divergence (KLD) metric. The blue boxplots measure divergence between each primary tumor type to each other metastatic tumor type, while the red boxplots measure divergence between each primary tumor type to every other primary tumor type. (C) Plotting the KLD of each tumor type’s primary-metastatic divergence with respect to the model’s performance as measured by the area under the receiver operator characteristics curve (AUROC) reveals weak positive correlation, suggesting that increases in the divergence between primary and metastatic tile populations may only marginally improve the efficacy of transfer learning.

We quantify the degree to which two non-overlapping distributions differ using the Kullback-Leibler divergence metric. This metric is computed for each of the learned latent features and visualized as a pair of boxplot distributions in Figure 5B which capture the difference between each class of primary tumors with respect to everything else in the dataset (red boxes), and with respect to the class’ metastatic counterparts (blue boxes). We compute the mean KL divergence across each of the learned features to quantify the statistical dissimilarity between primary and metastatic tiles across each of the fourteen classes, and then measure the correlation between mean KL divergence and the model’s reported per-class AUROC performance, shown in Figure 5C. We measure a slight linear association with a correlation of 0.317, suggesting a very slight association between a greater divergence between primary and metastatic samples and model’s classification performance in the transfer learning setting.

## 4 Discussion

This work illustrates that incorporating primary tumor histology into pre-training a histopathological classification of metastatic samples confers advantage in classification performance relative to models trained only on images of primary or only on images of metastatic tumors. Further, we illustrate that the degree to which primary and metastatic cancers may be statistically similar based on image content may affect the degree to which transfer learning benefits the modeling process.

This work has a number of limitations that may limit the extensibility of its findings. In particular, this study was limited to fourteen distinct classes of metastatic origin that were treated independently within the learning model. Future efforts may necessitate larger data sets with greater class-specific coverage to ensure robust ability to generalize both inter- and intra-class accuracy. This work also ignores shared latent features of tumor tissue that may be clinically relevant to rendering an accurate diagnosis of metastatic origin. Namely, this work ignores any other clinical feature of interest that may be relevant to this task, such as age, gender, medical history, and incidence of other disease phenotypes. This work also presents the clinical challenge of inferring metastatic origin in a simplified setting, when in practice a pathologist uses both other clinical covariate factors as well as obtainable results from axillary testing such as immunohistochemistry or genomic sequencing to guide their diagnostics.

The use of unsupervised feature extraction methods to infer feature divergence between primary and metastatic cancers also has a number of limitations. Like other efforts that incorporate these kinds of learning models, the learned feature space is subject to discrepancies, a lack of interpretability, and inconsistency in feature space embedding with similar input images. Although exploratory, it might be reasonable to use the relative dispersal of samples within a learned feature space to infer the heterogeneity or variability of intra-class samples so as to guide researchers in determining an adequate number of representative samples so as to cover an inferred feature space.

This work presents a number of future directions, in particular opening up the opportunity to explore clinical application of augmented inference of metastatic origin on a pathologist’s classification performance and with respect to the selection of confirmatory immunohistochemistry stains chosen to infer metastatic origin. Overall these results reinforce the importance of pre-training computer vision systems in digital pathology as a mechanism to overcome limitations in data set size for niche questions of narrow scope. Future efforts are expected to expand upon these findings by incorporating large public data sets into training paradigms to further enhance the capabilities of computer vision systems to infer the origin of metastatic cancers from whole slide histology.

## Acknowledgments

We extend our thanks to the staff at the OHSU Knight BioLibrary for their support in data access and dissemination. Further, we gratefully acknowledge the resources of the Exacloud high performance computing environment developed jointly by OHSU and Intel and the technical support of the OHSU Advanced Computing Center.

## Ethics approval and consent to participate

This study did not require ethical approval.

## Author contributions

GFS, EAB, CC, and YHC conceived of and developed the design of this study. CC and HG provided pathological annotation of images used throughout this work. GT and JWG supervised and provided direct technical contributions to this work.

## Data availability

The datasets generated and/or analyzed during the current study are not publicly available but are available from the corresponding author on reasonable request.

